# Differential vulnerability of neuronal subpopulations of the subiculum in a mouse model for mesial temporal lobe epilepsy

**DOI:** 10.1101/2022.06.02.494518

**Authors:** Julia Franz, Nicole Barheier, Susanne Tulke, Carola A. Haas, Ute Häussler

**Affiliations:** Experimental Epilepsy Research, Department of Neurosurgery, Medical Center - University of Freiburg, Faculty of Medicine, 79106 Freiburg, Germany; Faculty of Biology, University of Freiburg, 79104 Freiburg; Center for Basics in NeuroModulation, Faculty of Medicine, University of Freiburg, 79106 Freiburg, Germany; BrainLinks-BrainTools, University of Freiburg, 79110 Freiburg, Germany

**Keywords:** Parvalbumin, Neuropeptide Y, Calbindin, Calretinin, FluoroJade C, intrahippocampal kainate, mesial temporal lobe epilepsy, dorsoventral axis

## Abstract

Selective loss of inhibitory interneurons (INs) that promotes a shift toward an excitatory predominance may have a critical impact on the generation of epileptic activity. While research on mesial temporal lobe epilepsy (MTLE) has mostly focused on hippocampal changes, including IN loss, the subiculum as the major output region of the hippocampal formation has received comparatively little attention. Although it has been shown to occupy a key position in the epileptic network, data on cellular changes in the subiculum are controversial. Using the intrahippocampal kainate (KA) mouse model for MTLE, which recapitulates main features of human MTLE such as hippocampal sclerosis and granule cell dispersion following *status epilepticus* (SE), we identified cell loss in the subiculum and quantified changes in specific IN subpopulations along its dorsoventral axis.

We performed intrahippocampal recordings, FluoroJade C-staining for degenerating neurons shortly after SE and immunohistochemistry for neuronal nuclei (NeuN), parvalbumin (PV), neuropeptide Y (NPY), calretinin (CR) and calbindin (CB), and *in situ* hybridization for glutamic acid decarboxylase *(Gad) 67* mRNA at 21 days after KA.

We observed remarkable cell loss in the ipsilateral subiculum shortly after SE which was reflected in lowered density of NeuN+ cells in the chronic stage when epileptic activity could be measured in the subiculum concomitant with the hippocampus. We show a position-dependent reduction of *Gad67*-expressing INs by ~50% which particularly affected the PV- and to a lesser extent CR-expressing INs, whereas CB-expressing cells were unchanged.

Interestingly, the density of NPY-positive neurons was increased, but double-labeling for *Gad67* mRNA expression revealed that rather a *de novo* expression of NPY in non-GABAergic cells instead of IN changes underlay this increase.

Our data reveal a position- and cell type-specific vulnerability of subicular INs in MTLE which might contribute to hyperexcitability of the subiculum, as reflected in the occurrence of epileptic activity.

**Keypoints (3-5):**

- The subiculum develops epileptic activity after intrahippocampal kainate injection in mice
- *Gad67*-mRNA expressing interneurons are reduced in the subiculum in the intrahippocampal kainate model for mesial temporal lobe epilepsy
- Parvalbumin- and calretinin-expressing interneurons are particularly vulnerable in epilepsy
- Neuropeptide Y upregulation in non-GABAergic cells in the subiculum indicates compensatory processes

## Introduction

Mesial temporal lobe epilepsy (MTLE) is among the most frequent forms of pharmacoresistant epilepsies. To elucidate the underlying pathophysiological mechanisms, the hippocampus as site of origin of focal seizures has been intensely investigated. This revealed histological alterations including hippocampal sclerosis, granule cell dispersion and mossy fiber sprouting as hallmarks of MTLE in patients and animal models^1–8^. In contrast, the subiculum received less attention, despite playing a pivotal role in the network as a major output region of the hippocampal formation.

Indeed, there is evidence that the subiculum does not undergo relevant cell loss in MTLE^9,10^, but studies in animal models for MTLE challenge this assumption by showing decreased neuronal density^11,12^. Moreover, the subiculum becomes subject of considerable deafferentation due to extensive cell loss in the CA1 region in patients^5^ and animal models^13^. With its large proportion of bursting cells^14,15^ and a network that facilitates synchronization of neuronal activity^16,17^, the subiculum is an ideal candidate region for the generation of epileptic activity. Using brain slices from MTLE patients, it has been proposed that the subiculum and not the hippocampus is the origin of spontaneous interictal activity^18,19^, which was confirmed by *in vivo* recordings in the subiculum of patients with pharmacoresistant MTLE^20^. This activity is suggested to arise from impaired inhibition by GABAergic interneurons (INs)^18^. In the hippocampus, IN subpopulations were shown to be differentially vulnerable to epileptic activity^21–23^, whereas less is known about vulnerability of subicular INs.

We used the intrahippocampal kainate (ihKA) mouse model which displays recurrent epileptic activity and major histological hallmarks of MTLE^7,24^. Despite the focal injection, structural changes and epileptic activity are not limited to the injection site but involve different hippocampal subregions, large parts of the dorsoventral (DV) axis, and both hippocampi to different degrees^23,25,26^. Therefore, we investigated whether IN subpopulations are altered in the subiculum along the DV axis in both hemispheres to provide an additional source of impaired excitation-inhibition balance. We measured the occurrence of epileptic activity in the subiculum and detected degenerating neurons in the ipsilateral subiculum already 48h after ihKA, and NeuN staining substantiated reduced neuronal density in chronic epilepsy. Furthermore, our quantitative analysis revealed position- and cell type-specific vulnerability of IN populations in the subiculum under epileptic conditions.

## Materials & Methods

### Animals

Experiments were performed in adult (>8 weeks) C57Bl/6 wildtype mice or negative littermates of transgenic lines bred on C57Bl/6 background (CEMT Freiburg). Mice were housed under standard conditions (12h-light/dark, room temperature (RT), food and water *ad libitum).* Experiments were carried out following the guidelines of the European Community’s Council Directive of 22 September 2010 (2010/63/EU), approved by the regional council (Regierungspräsidium Freiburg). In total, 46 mice were used (35 male, 11 female; 16 NaCl, 30 KA).

### Surgery and in vivo recordings

Surgery was performed under deep anesthesia (ketamine hydrochloride 100mg/kg body weight, xylazine 5mg/kg, atropine 0.1mg/kg, i.p.) with analgesic treatment (buprenorphin hydrochloride 0.1mg/kg, carprofen 5mg/kg, s.c.; lidocaine 2mg/kg, s.c. to scalp). Mice were mounted in a stereotaxic frame and injections of 50nL KA (20mM in 0.9% NaCl, Tocris) or 0.9% NaCl-solution into the right dorsal hippocampus were performed using a Nanoject III (Drummond Scientific Company) at coordinates anterioposterior [AP]=-2.0mm, mediolateral [ML]=-1.4mm and DV=-1.8mm from bregma. In KA-injected mice, behavioral *status epilepticus* (SE) with head nodding, convulsions, rotations and immobility was verified as previously described^27^. Postsurgical analgesia was performed with carprofen for 2 days.

One week after ihKA/ihNaCl injection, a subgroup of mice (6 NaCl, 9 KA) were implanted with wire electrodes (platinum-iridum, teflon-insulated, Ø125μm, WPI) into the dentate gyrus (DG; AP=-2.0mm, ML=±1.4mm, DV=-1.9mm) and the subiculum (AP=-3.0mm, ML=±1.4mm, DV=-1.4mm) of both hemispheres (anesthesia see above, analgesia with burpenorphin+carprofen for 2 days). Jewelers’ screws above the frontal cortex served as ground/reference. A connector was permanently fixed to the skull with cyanoacrylate and dental cement. Freely moving mice were recorded (4-6 sessions, ~2h each) between 14 and 28 days after ihKA connected to a miniature preamplifier (Multi Channel Systems). Signals were amplified (500-fold, band-pass filter 1Hz-5kHz) and digitized (sampling rate 10kHz, Spike2 software; CED). Mice with misplaced electrodes verified by posthoc histology or bad connections were excluded (3 NaCl, 3 KA), two ihKA mice died after implantation.

### Perfusion and slice preparation

Mice were transcardially perfused under deep anesthesia with 0.9% saline followed by paraformaldehyde (PFA, 4% in 0.1M phosphate buffer (PB), pH 7.4). Brains were post-fixed in PFA overnight (4°C), cryoprotected (30% sucrose) and frozen in isopentane. Horizontal sections (50μm) were prepared on a cryostat and collected in PB for immunohistochemistry or in 2x saline sodium citrate (SSC) for fluorescence *in situ* hybridization (FISH).

### FluoroJade C

FluoroJade C (FJC, Merck Millipore) staining was used to monitor cell death 2 and 4 days after ihKA injection. Sections were mounted on gelatine-coated slides and dried overnight (RT), re-hydrated (1% NaOH in 100% Ethanol for 5min; 70% Ethanol for 2min, distilled water for 2min), incubated in 0.06% potassium permanganate (10min), rinsed and transferred in 0.0001% FJC solution (10min, darkness), followed by washing, xylene immersion and coverslipping with Hypermount.

### Immunohistochemistry

After preincubation with 0.25% Triton X-100 and 10% normal horse/goat serum in 0.1M PB (30min, free-floating), sections were incubated overnight (4°C) with primary antibodies: mouse anti-CB (1:2000; #300), rabbit anti-calretinin (CR; 1:500; #7699/4), rabbit anti-parvalbumin (PV; 1:1000; #PV27, all Swant), guinea pig anti-NeuN (1:500; #266004, Synaptic Systems) and rabbit anti-NPY (1:1000; #ab30914, Abcam). Secondary antibodies, marked with Cy2, Cy5 (both 1:200; Jackson ImmunoResearch) or Alexa555 (1:500; Thermo Fisher Scientific) were applied (2-3h, RT). A mouse-on-mouse kit (MOM Basic Kit; Vector Laboratories) was used for CB. Sections were counterstained with 4’,6-diamidino-2-phenylindole (DAPI, 1:10,000). When combining FISH and immunohistochemistry, Triton X-100 was omitted. Sections were coverslipped with fluorescence mounting medium (Immu-Mount, Thermo Shandon Ltd or Dako).

### Fluorescence in situ hybridization

Expression of *Gad67* mRNA was investigated by FISH with digoxigenin (DIG)-labeled cRNA probes. Probes were generated by *in vitro* transcription from appropriate plasmids^28^. Sections were hybridized overnight at 45°C, followed by detection with an anti-DIG antibody and tyramide signal amplification as described^23,27^.

### Microscopy and quantification

Sections were imaged with an epifluorescence microscope (AxioImager 2, Zen software, Zeiss; 10-fold Plan Apochromat objective, NA=0.45; AxioCam 506). Cell counting was performed with ImageJ (version 1.52a; NIH) on all available brain sections from DV −1,2 to - 3,2mm relative to bregma. The region of interest (ROI, principal cell layers of the subiculum) was drawn according to a brain atlas^29^. Cells were counted manually using the cell counter plugin by a single experimenter for consistency. Criteria for selection were: signal intensity clearly distinguishable from background, cell shape and presence of a DAPI-stained nucleus. Blinding to the treatment was impossible due to KA-induced alterations in the hippocampus. To compare control and KA-injected mice along the DV axis, sections were assigned the following subgroups: dorsal to the injection site (−1,2 to −1,8mm – usually 3 sections/mouse), approximately at the injection site (−1,9 to −2,1mm – 2 sections/mouse) and ventral (−2,2 to maximally −3,2mm – 4-6 sections/mouse). Cell density was calculated for the ROI (cells/mm^2^) and averaged within subgroups per animal. Values for the ipsi- and contralateral subiculum of NaCl-injected mice were grouped since we did not observe any differences.

### Statistics

Statistical analysis was performed with GraphPad Prism (version 9.3.1). A Shapiro-Wilk test was performed to test for normality. In case of normality a one-way analysis of variance (ANOVA) was followed by Tukey’s multiple comparisons test, otherwise a Kruskal-Wallis test was followed by Dunn’s multiple comparisons test.

## Results

### Epileptic activity occurs in the dentate gyrus and in the subiculum

In controls (n=3), LFP patterns during immobility consisted of irregular activity and occasional dentate spikes in the DG and sharp-wave ripples in the subiculum (Fig. 1A). In ihKA mice (n=4) we recorded recurrent epileptiform activity in the DG, but also in the subiculum (Fig. 1B). Fast ripples riding on large population spikes were evident in the DG (Fig. 1B1) and in the subiculum where they resembled sharp-wave-ripples (SWRs, Fig. 1B2). In 3/4 mice generalized convulsive seizures occurred during the 2h sessions which not only involved the ipsilateral DG and subiculum, but also the contralateral side where large epileptiform population spikes were measured (Fig. 1C,C1). In agreement with previous observations^30^, it was impossible to determine the exact onset site whereas the post-ictal depression started with a delay on the contralateral side (Fig. 1C2).

**Figure 1:**
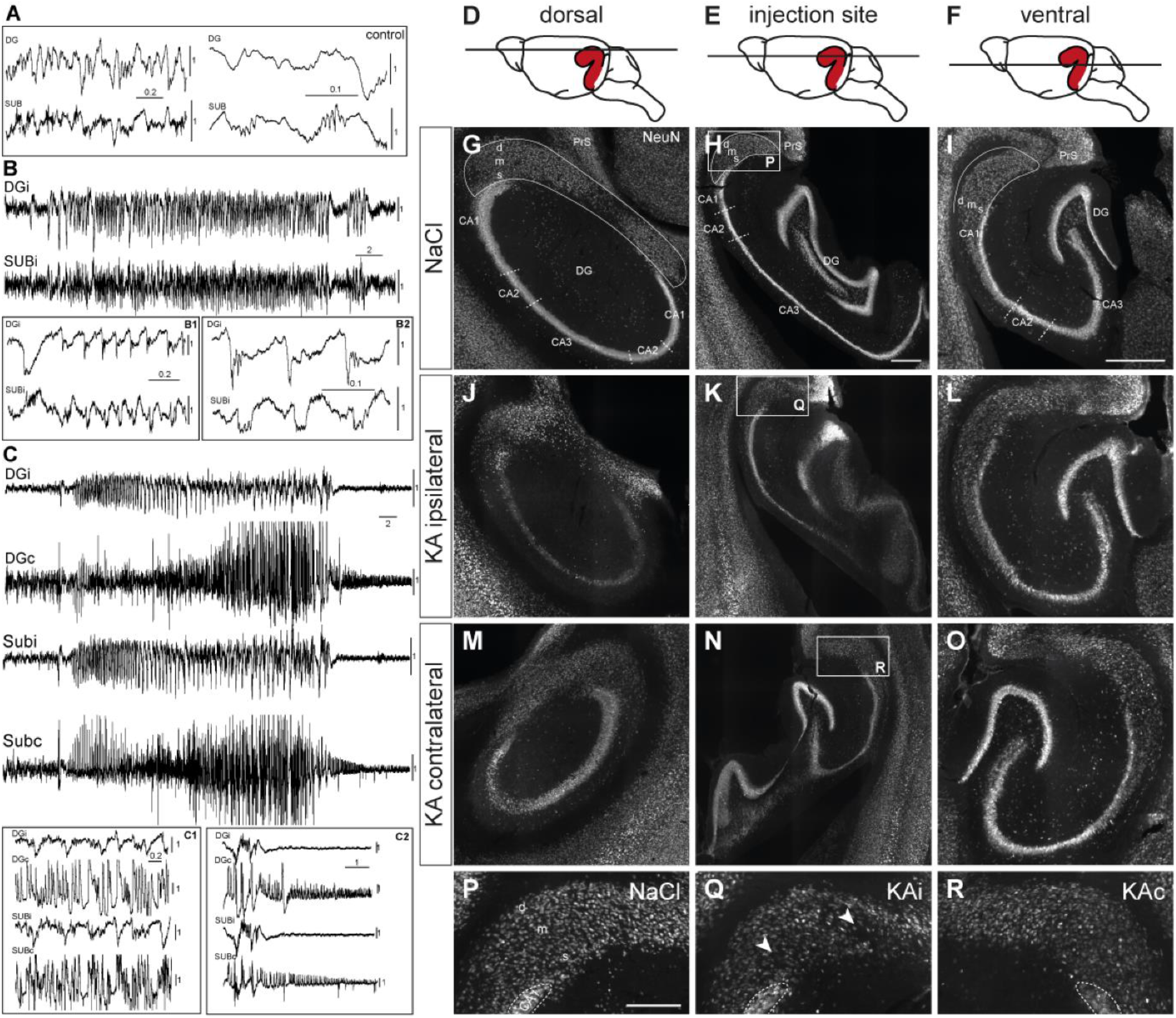
Epileptiform activity and neuronal loss in the subiculum. **A)** Local field potential (LFP) recording of a control mouse during awake immobility shows irregular activity in the dentate gyrus (DG) and subiculum (SUB) with sharp-wave-ripple complexes in the subiculum (enlarged in traces at the right side). **B)** KA-injected mouse. Subclinical epileptic seizure (during immobility, without behavioral correlate) involved the ipsilateral DG (DGi) and subiculum (SUBi). Large epileptiform populations spikes are enlarged in B1 and B2, B2 also shows high frequency oscillations riding on population spikes in the DG and the subiculum. **C)** Generalized convulsive seizure recorded in the DG and subiculum of both hemispheres. C1 displays a cutout depicting the large amplitudes on the contralateral side. Seizures are followed by post-ical depression (C2) which starts earlier on the ipsilateral than on the contralateral side. Scale bars in A-C: horizontal – time in s, vertical – voltage in mV. Channels are individually scaled for better visibility of characteristic patterns. **D-F)** Schematic drawings represent the three positions (**D** dorsal, **E** injection site, **F** ventral) that were investigated along the dorsoventral (DV) axis. **G-R)** NeuN immunostaining. Horizontal sections of the hippocampal formation from NaCl-injected control mice (**G-I**) were compared to 21 days after kainate (KA, **J-R**). The position of the subiculum is encircled in G-I. **J**) Ipsilateral side, dorsal position. At 21 days after KA the characteristic extensive cell loss in the CA regions and granule cell dispersion confirms successful KA injection. Cell density in the subiculum is reduced. **K**) Ipsilateral side, DV level of the injection site. Cell loss in the CA region and subiculum together with dispersion of the granule cell layer is visible. **L**) Ipsilateral side, ventral position. At this DV level, the hippocampus and subiculum appear similar to control concerning density and localization of NeuN-positive cells. **M-O)** Neuronal densities of the contralateral hippocampus and subiculum are comparable to controls at all DV levels. **P-R)** Cutouts (white box in H,K,N) illustrate the subiculum at the level of the injection site. **P)** Dense arrangement of neurons in the subiculum after NaCl injection. The deep (d), middle (m) and superficial (s) layer of the subiculum is indicated. **Q)** Decreased neuronal density in the ipsilateral subiculum under epileptic conditions. Arrows mark positions of pronounced cell loss in the proximal subiculum close to CA1 and in the distal subiculum close to the border to the presubiculum. **R)** The contralateral subiculum after KA injection is comparable to control, but NeuN staining is weaker in some mice. Scale bars: D-L 500 μm, M-O 250 μm. CA *cornu ammonis*, DG dentate gyrus, d deep pyramidal cell layer, m middle pyramidal cell layer, s superficial pyramidal cell layer.

### Position- and layer-specific neuronal loss in the subiculum

To determine whether the subiculum is structurally altered in chronic MTLE, we performed immunohistochemistry for NeuN at 21 days after ihKA (Fig. 1, n=8). We will refer to the part of the subiculum adjoining CA1 as proximal and the area adjacent to the presubiculum as distal and distinguish superficial, middle and deep layers with the superficial layer adjoining the almost cell-free molecular layer, according to previous work^31^. We compared three levels along the DV axis in horizontal sections: dorsal to the injection site (Fig. 1D), around the injection site (Fig. 1E) and ventral (Fig. 1F).

Extensive cell loss in areas CA1 and CA3 of the hippocampus at the dorsal and the injection site level, accompanied by granule cell dispersion at the level of the injection site confirmed the successful KA injection (Fig. 1J,K). Regarding the dorsal subiculum, reduced density of NeuN^+^ cells was visible in proximal and distal parts, mainly in the superficial and middle layer, in comparison to controls (Fig. 1G,J). At the level of the injection site NeuN^+^ cells were also thinned out throughout the ipsilateral subiculum (Fig. 1H,K), mainly in the superficial and middle layer (Fig. 1Q). Moreover, many of the remaining neurons in the subiculum in both hemispheres displayed weaker NeuN staining than controls (Fig. 1P-R). The latter was also visible in the ventral subiculum, yet without any clear sign of cell loss (Fig. 1I,L). In the contralateral subiculum neuronal density was comparable to controls at all levels (Fig. 1M-O). To determine whether the reduced density is due to SE-induced neurodegeneration, we performed FJC staining at 2 and 4 days after ihKA injection. At 2 days, degenerating neurons were densely arranged in ipsilateral CA1 (Fig. 2A,B) and all over the ipsilateral hippocampus (data not shown), as described previously^23^. In the subiculum, FJC^+^ cells were more abundant throughout all layers at the dorsal position compared to the injection site and the ventral position, where they were mainly located in superficial and middle layers (Fig. 2A-C, n=4). At 4 days after ihKA, FJC^+^ cells showed a similar distribution, but with overall slightly lower density (Fig. 2D-F, n=6). Contralaterally, there were only very few FJC^+^ cells both in the hippocampus and the subiculum at all levels (Fig. 2G-I, 2 days not shown).

**Figure 2:**
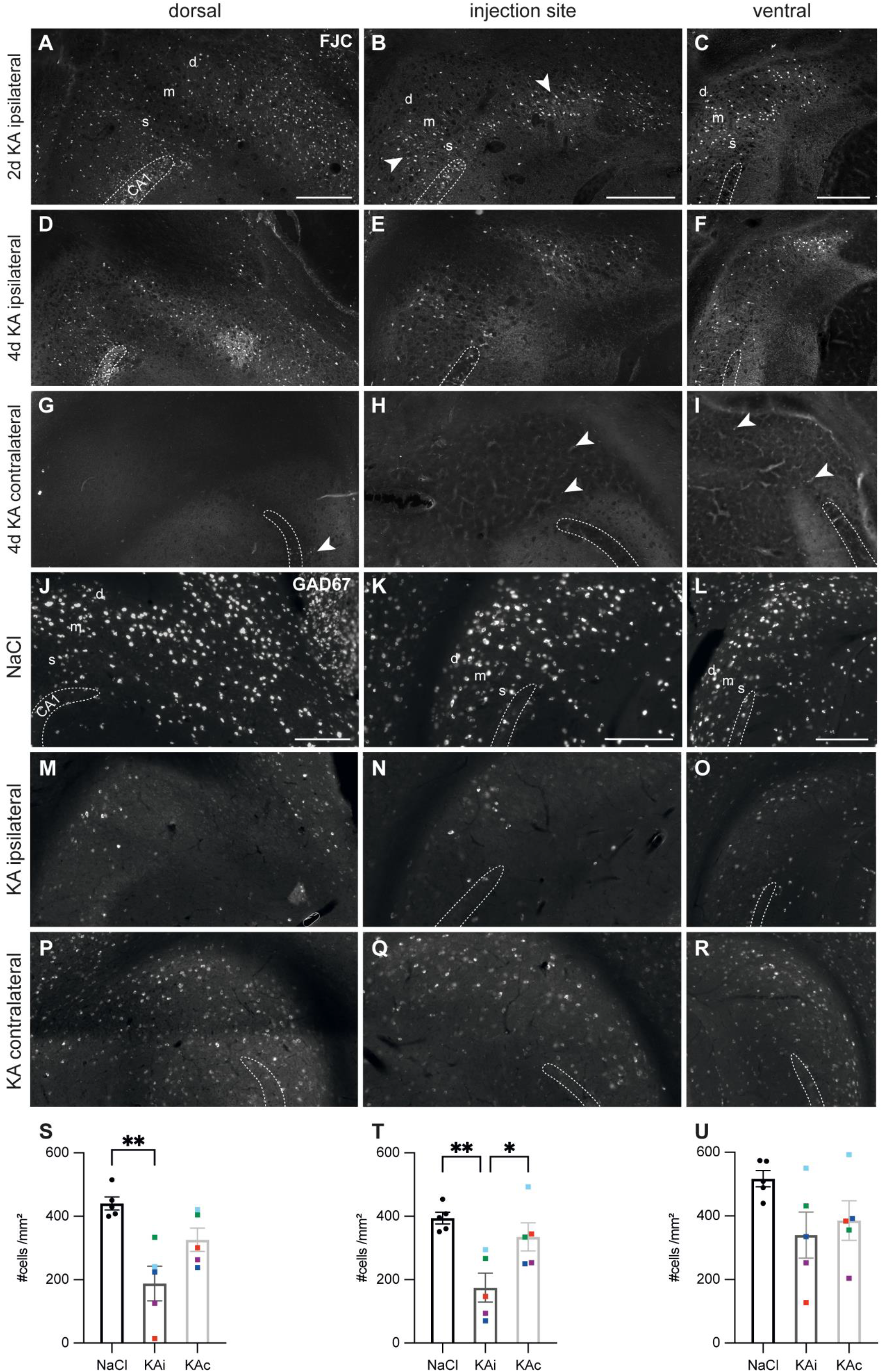
Degenerating neurons in the ipsilateral subiculum early after kainate injection and strong loss of GABAergic interneurons in the chronic stage. **A-I)** FluoroJade C (FJC) staining was performed to mark degenerating neurons at 2 days (**A-C**) and 4 days after KA injection (**D-I**) along the DV axis. Representative sections are shown, borders of the CA1 pyramidal cell layer are indicated. **A)** At 2 days after KA, FJC^+^ cells are distributed throughout the pyramidal cell layer (superficial (s), middle (m), and deep (d) layer) of the dorsal ipsilateral subiculum. **B)** At the level of the injection site, FJC-stained neurons are mainly located in the superficial and middle pyramidal cell layer. Degenerating neurons are densely placed in the proximal subiculum close to CA1 and in the distal subiculum at the border to the presubiculum (arrows). **C)** Similar pattern in the ventral subiculum but with lower density of FJC-positive cells. **D-F)** At 4 days after KA the distribution of FJC^+^ cells is similar to 2d, but less dense at all DV positions. **G-I)** In the contralateral subiculum, only a few neurons were marked along the DV axis. **J-R)** Representative horizontal sections of a fluorescent *in situ* hybridization for *Gad67* mRNA in a control (**J-L**) and an epileptic (**M-R**) mouse 21 days after kainate (KA) injection. **J)** Dorsal subiculum after NaCl injection. GABAergic interneurons are equally distributed in all layers without a difference between the proximal and distal subiculum. **K)** At the level of the injection site, the number of *Gad67* mRNA expressing cells is similar to the dorsal position. **L)** Ventrally the density of GABAergic interneurons is slightly higher. **M-O)** Ipsilateral subiculum 21 days after KA injection, dorsal position (**M**) and injection site (**N**). The density of *Gad67* mRNA expressing cells is strongly decreased. **O)** At the ventral position more *Gad67* mRNA-expressing cells were preserved than at the more dorsal sites. **P-R)** Contralateral subiculum 21 days after KA injection. The density of *Gad67* mRNA expressing neurons is comparable to control at all DV levels but *Gad67* mRNA labeling was weaker. **S-U)** Quantification of *Gad67* mRNA expressing cells in the subiculum of NaCl-injected mice (ipsi- and contralateral side combined), ipsi- and contralateral subiculum of KA-injected mice 21 days after injection. Individual animals are color-coded, bars show means (cells/mm^2^) with standard error of the mean (SEM). **S)** In the dorsal subiculum *Gad67* mRNA-expressing cells in the ipsilateral subiculum 21 days after KA injection are significantly reduced compared to controls (one-way ANOVA: p=0.003; Tukey’s multiple comparisons test: NaCl-KAi p=0.002). **T)** At the level of the injection site the density of *Gad67* mRNA-expressing cells in the ipsilateral subiculum is significantly reduced compared to the contralateral subiculum and controls (one-way ANOVA: p=0.004; Tukey’s multiple comparisons test: NaCl-KAi p=0.004, KAc-KAi p=0.029). **U)** At the ventral position, density of GABAergic interneurons varies strongly across epileptic mice, means are not different (one-way ANOVA p=0.115). Scale bars: 250 μm. *p<0.05, **p<0.01. CA1 *cornu ammonis* 1, d deep pyramidal cell layer, m middle pyramidal cell layer, s superficial pyramidal cell layer.

Together, early cell loss and reduced neuronal density in the chronic stage confirm ihKA-induced structural changes in the subiculum, yet less prominent than in the hippocampus. Next, we focused on selected IN populations.

### Loss of GABAergic interneurons in the subiculum

In controls, *Gad67* mRNA-expressing INs were densely arranged throughout all cell layers of the subiculum (Fig. 2J-L, n=5). Quantitative analysis did not reveal any major differences between the proximal and the distal subiculum, as well as between the dorsal subiculum and the level of the injection site (Fig. 2J-L,S,T). In the ventral subiculum, the density of *Gad67* expressing cells was slightly higher (Fig. 2U).

At 21 days after ihKA (n=5), *Gad67* mRNA^+^ INs were strongly diminished throughout the dorsal ipsilateral subiculum (Fig. 2M) and at the level of the injection site (Fig. 2N), and in some mice they still appeared less dense further ventrally (Fig. 2O). In addition, the *Gad67* FISH was less intense indicating reduced mRNA levels per cell. In the contralateral subiculum, the density of *Gad67*^+^ cells was comparable to control but the staining intensity was also lower at all levels (Fig. 2P-R). Quantification revealed the loss of ~50% of *Gad67^+^* INs in the dorsal ipsilateral subiculum and at the level of the injection site (Fig. 2S,T). Ventrally, mean values did not differ from controls and individuals ranged from a strong loss to normal density (Fig. 2U), neither did the density of *Gad67^+^* INs in the contralateral subiculum differ from controls.

### Loss of parvalbumin-positive interneurons in the ipsilateral subiculum

In control mice (n=5), INs expressing the Ca^2+^-binding protein PV were scattered throughout all cell layers of the subiculum with their fibers building a honeycomb-like pattern reaching into the molecular layer (Fig. 3A-C). Their density was comparable at all DV positions (Fig. 3J-L). At 21 days after ihKA (n=5), we observed a reduction of PV^+^ INs at all DV positions of the ipsilateral subiculum (Fig. 3D-F), but most pronounced dorsally (Fig. 3D). At the injection site and in the ventral subiculum, the superficial and middle layers were more affected than the deep layer (Fig. 3E,F). Some of the remaining PV^+^ cells appeared hypertrophic (Fig. 3E). Quantitative analysis confirmed a significant reduction of PV^+^ INs in the dorsal ipsilateral subiculum by ~43% (Fig. 3J). The lower mean density at the injection site and in the ventral subiculum was not significant (Fig. 3K,L). On the contralateral side, the distribution and density of PV^+^ cells in the dorsal and ventral subiculum was comparable to controls (Fig. 3G,I,J,L), whereas density tended to be lower at the level of the injection site (Fig. 3H,K). In two mice the signal in the dorsal contralateral subiculum appeared stronger, possibly due to stronger PV expression in somata and axons and/or increased ramification of processes (Fig. 3G).

**Figure 3:**
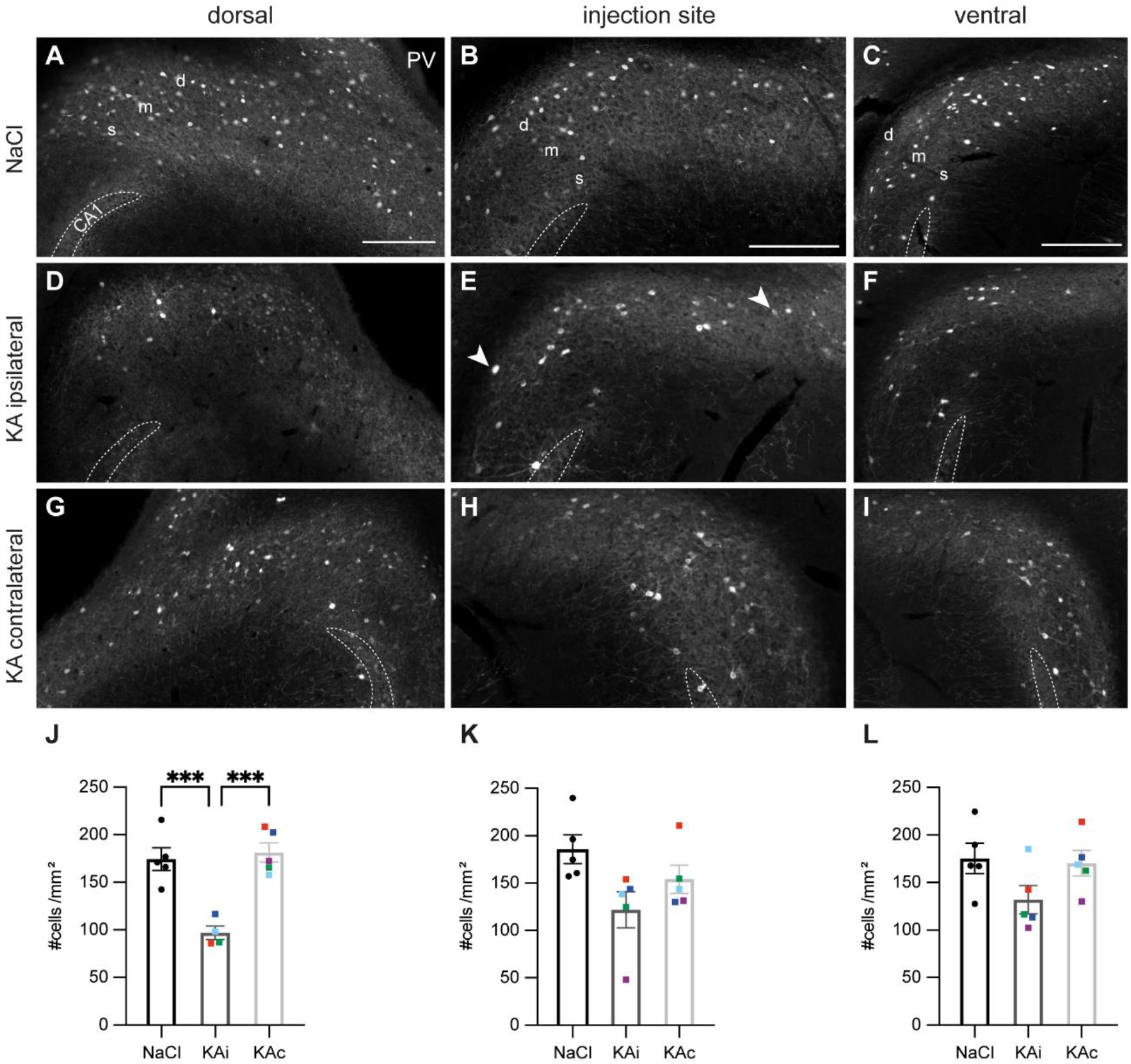
Parvalbumin-positive interneurons are reduced in the ipsilateral subiculum 21 days after KA. Representative sections immunostained for parvalbumin (PV) in control mice (**A-C**) and 21 days after kainate (KA) injection (**D-I**). Borders of the CA1 pyramidal cell layer are indicated in all panels. **A)** Dorsal subiculum after NaCl injection. PV^+^ cells are evenly distributed throughout all layers of the proximal and distal subiculum (s superficial, m middle, d deep) and PV^+^ fibers form a dense plexus. **B, C)** A comparable pattern is visible at the level of the injection site (**B**) and further ventral (**C**). **D)** Ipsilateral subiculum 21 days after KA, dorsal position. PV-expressing somata and PV^+^ fibers are strongly diminished, mainly in the superficial and middle layers. **E**) Similar pattern for the level of the injection site and further ventral (**F**). Some PV^+^ interneurons appear hypertrophic (examples are marked with arrows in **E**). **G-I**) Contralateral subiculum 21 days after KA. At the dorsal (**G**) and ventral (**I**) position, density of PV-expressing interneurons appears unchanged. **H)** A slight reduction of PV^+^ cells occurred at the level of the injection site. **J-L)** Quantification of PV expressing cells in the subiculum of NaCl-injected mice, ipsi- and contralateral subiculum of KA-injected mice 21 days after injection. Individual animals are color-coded, bars show means (cells/mm^2^) with standard error of the mean (SEM). **J)** In the dorsal ipsilateral subiculum (KAi) the density of PV-expressing cells is reduced by more than 40%, whereas the contralateral subiculum (KAc) does not differ from controls (NaCl; one-way ANOVA: p<0.001, T ukey’s multiple comparison test NaCl – KAi: p<0.001; KAi – KAc: p<0.001). **K)** The lower mean density in the ipsilateral subiculum at the injection site was not significantly different from the contralateral subiculum or control (one-way ANOVA: p = 0.05). **L)** The effect of KA on PV-expressing interneurons is weaker in the ventral subiculum (one-way ANOVA: p = 0.12). Scale bars: 250 μm. ***p<0.001. CA1 *cornu ammonis* 1, d deep pyramidal cell layer, m middle pyramidal cell layer, s superficial pyramidal cell layer.

### Reduction of neuropeptide Y-expressing interneurons but compensatory NPY upregulation

In dorsal sections of control mice (n=5), NPY^+^ neurons were scattered in superficial to deep layers of the proximal and distal subiculum (Fig. 4A). Further ventral, NPY^+^ neurons were primarily found in the proximal subiculum with intense arborizations in the molecular layer (Fig. 4B,C).

**Figure 4:**
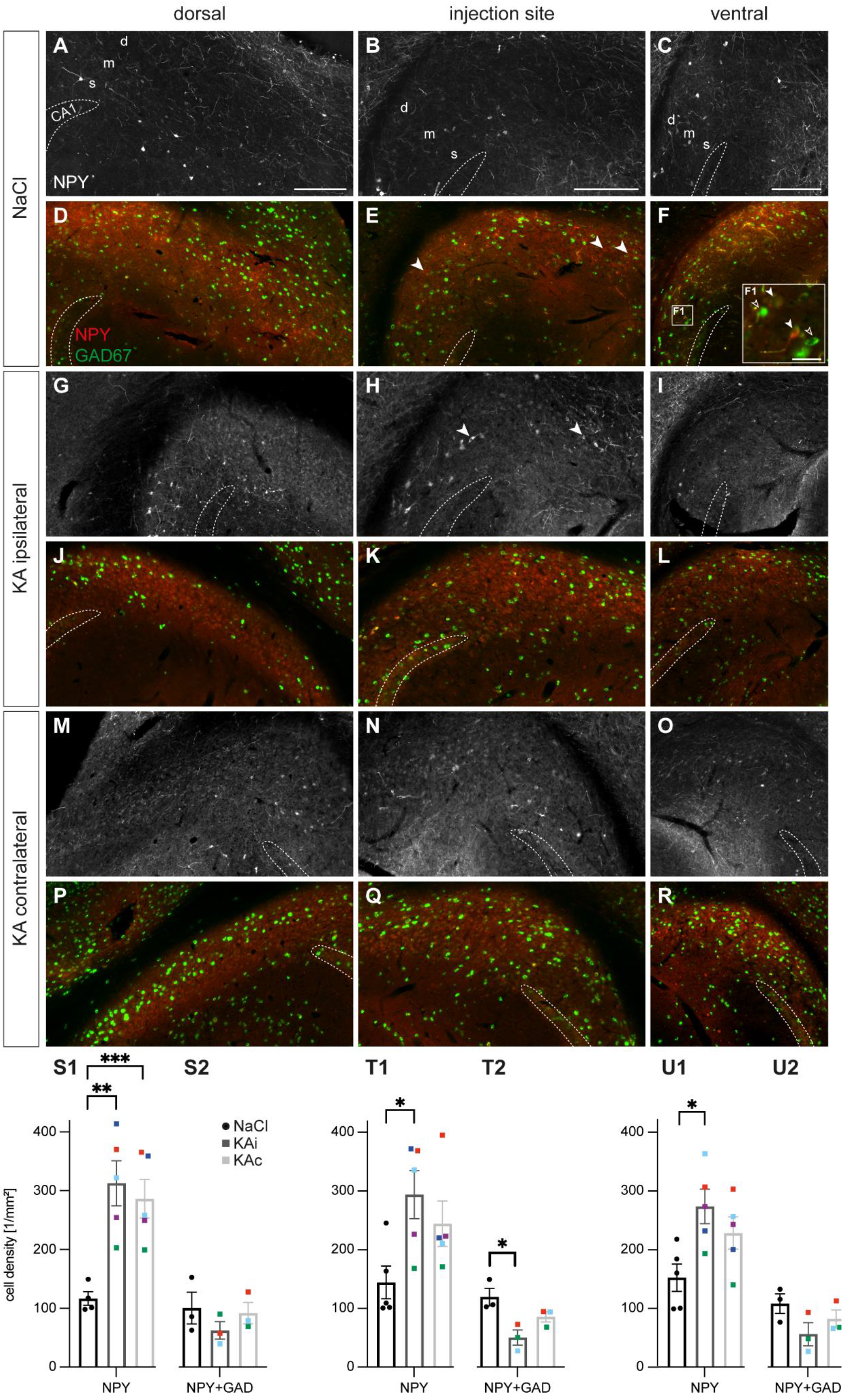
Kainate induces an upregulation of neuropeptide Y, but causes the loss of neuropeptide Y interneurons. Representative images depicting the subiculum of control (**A-F**) and kainate (KA)-injected mice (**G-R**). Immunolabeling of Neuropeptide Y (NPY)-expressing cells (red, **A-C**, **G-I**, **M-O**) and counterstaining for co-expression of *Gad67* mRNA using fluorescent *in situ* hybridization (green, **D-F**, **J-L**, **P-R**). Borders of the CA1 pyramidal cell layer are indicated. **A)** NaCl-injected control, dorsal subiculum. NPY^+^ cells and their processes are located mainly in the superficial and deep layer of the proximal and distal subiculum. **B)** At the injection site labeled cells are more abundant in the proximal subiculum. **C)** Ventral subiculum, distribution of NPY^+^ neurons similar to (**B**). **D-F)** High density of *Gad67* mRNA-expressing neurons (green) at all positions along the DV axis in control mice. A subset of these cells is NPY-positive (overlay in yellow), whereas a few NPY-expressing cells are *Gad67*-negative (arrows in **E**). **F1)** Enlarged cutout marked in (**F**). Double-positive cells (filled arrows) and *Gad67* mRNA-positive but NPY-negative neurons (empty arrows) are visible. **G)** At 21 days after KA, NPY expression is strongly increased throughout the dorsal ipsilateral subiculum. **H, I)** Same for the injection site and the ventral position. Cells of various shapes and sizes are NPY-positive (arrows mark examples). **J)** The density of neurons expressing *Gad67* mRNA is decreased in the ipsilateral subiculum after KA. Only a subset is NPY-positive. **K, L)** At the injection site and further ventral, *Gad67* mRNA-expressing cells are decreased, but to a lesser extent. Only a subset is NPY-positive. **M-O)** In the contralateral subiculum of KA-injected mice, NPY expression is upregulated at all positions along the DV axis. **P-R)** The distribution of *Gad67* mRNA-expressing neurons and of double-labeled cells is comparable to the control condition. **S1-U1)** Quantification of NPY-expressing cells in controls and at 21 days after KA. Individual values (color-coded for each animal), means and standard error of the mean (SEM) are displayed. **S1)** At the dorsal position the number of NPY-positive cells is massively increased in the ipsi- and contralateral subiculum compared to control (one-way ANOVA, p = 0.003; Tukey’s multiple comparison test: NaCl-KAi p = 0.004, NaCl-KAc p = 0.009). **T1)** At the injection site, mean density of NPY-positive cells is higher than in controls, but with high variability across animals. No significant differences were measured (not normally distributed, Kruskal-Wallis-Test p=0.048, Dunn’s multiple comparison test not significant for all pairs). **U1**) Increase of NPY-expressing cells in the ventral ipsilateral subiculum (one-way ANOVA: p = 0.023; Tukey’s multiple comparisons test: NaCl-KAi p = 0.02). **S2-U2)** Quantification of the density NPY/*Gad67* double-labeled cells (i.e., NPY-positive interneurons) in the subiculum. A trend for lower values in the ipsilateral subiculum after KA was visible for all DV positions, but reached significance only at the injection site (S2: one-way ANOVA: p = 0.44, T2: one-way ANOVA: p = 0.02, Tukey’s multiple comparison test: NaCl-KAi p = 0.02; U2: one-way ANOVA: p = 0.18). Scale bars: A-R 250 μm, F1 50 μm. *p<0.05, **p<0.01, ***p<0.001. CA1 *cornu ammonis* 1, d deep pyramidal cell layer, m middle pyramidal cell layer, s superficial pyramidal cell layer.

At 21 days after ihKA (n=5), NPY staining was strongly increased, which was reflected in much higher density of NPY^+^ somata and axonal processes in the dorsal ipsilateral (Fig. 4G) and contralateral subiculum (Fig. 4M). NPY expression was also increased at the level of the injection site and ventrally, predominantly in superficial and middle layers of the distal ipsilateral subiculum (Fig. 4H, I) and slightly less on the contralateral side (Fig. 4N,O). Overall, NPY^+^ cells appeared very diverse in size, shape and staining intensity after KA injection. Quantitative analysis confirmed a significant increase in density of NPY^+^ cells along the entire DV axis of the ipsilateral subiculum, with almost threefold higher numbers at the dorsal position (Fig. 4S1) and a twofold rise in more ventral sections (Fig. 4T1,U1). On the contralateral side, density of NPY^+^ cells also showed an ~threefold increase in the dorsal subiculum (Fig. 4S1), and tended to be higher at the other positions (Fig. 4T1,U1).

Is there an increase of NPY-expressing INs in the subiculum? Previous studies on NPY-expressing cells in the epileptic hippocampus have reported a transient upregulation in excitatory cells alongside a decreased number of NPY^+^ INs^23^. We therefore combined NPY immunohistochemistry with FISH for *Gad67* mRNA in a subset of samples. Only a small fraction of *Gad67* mRNA^+^ cells expressed NPY in controls (n=3; Fig. 4D-F,F1). Interestingly, a few NPY^+^ cells did not express *Gad67* mRNA (Fig. 4E), which explains overall smaller numbers in the quantification of controls (Fig. 4S-U).

At 21 days after ihKA, a subset of the NPY^+^ cells co-expressed *Gad67* mRNA, whereas most were negative, indicating the upregulation of NPY in non-GABAergic cells (n=3, Fig. 4J-L). In the contralateral subiculum, the density and distribution of *Gad67*^+^/NPY^+^ neurons was comparable to control (Fig. 4P-R). Quantification of *Gad67*^+^/NPY^+^ cells revealed a tendency towards lower numbers of double-labeled cells for the ipsilateral subiculum, which was significant for the level of the injection site (Fig. 4S2-U2). Together our data show a loss of NPY^+^ INs and NPY upregulation in the subiculum in non-GABAergic cells.

### Loss of calretinin-, but not calbindin-expressing interneurons

Lastly, we analyzed the distribution of cells expressing the calcium-binding proteins, CR and CB. Cells expressing CR were sparsely scattered mainly in the superficial and middle layers of all DV levels in controls (n=5, Fig. 5A-C).

**Figure 5:**
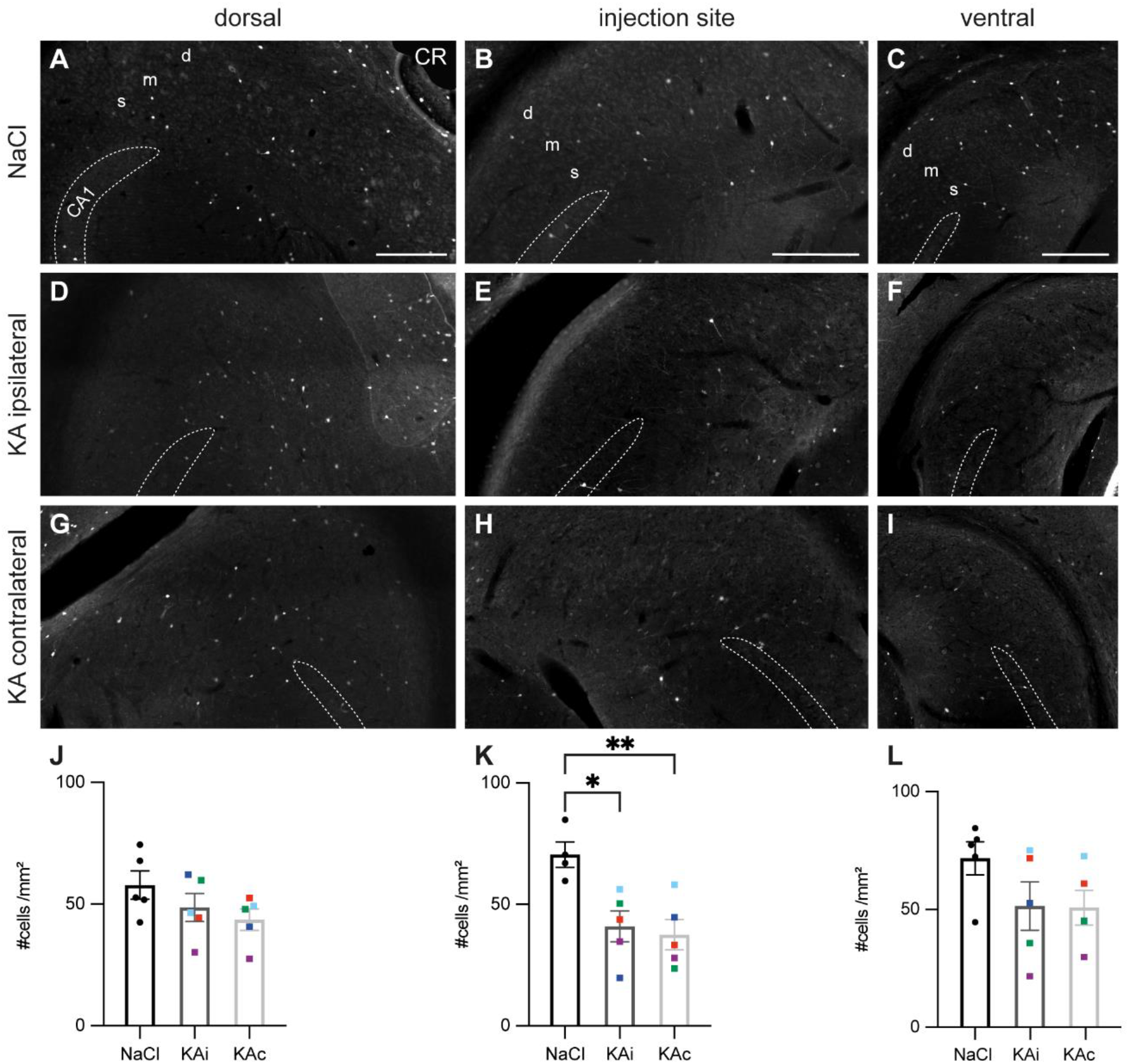
Calretinin-expressing neurons are reduced 21 days after kainate injection. Representative sections immunostained for calretinin (CR) in control mice (**A-C**) and 21 days after kainate (KA) injection (**D-I**). Borders of the CA1 pyramidal cell layer are indicated in all panels. **A-C)** In the subiculum of control animals CR-expressing neurons are sparsely scattered of CR throughout the pyramidal cell layer at the dorsal position (**A**), at the injection site (**B**) and more ventral (**C**). **D)** At 21 days after KA, the density and distribution of CR-positive cells is comparable to controls in the dorsal ipsilateral subiculum. **E)** At the injection site CR^+^ cells are reduced mainly in the proximal subiculum. **F)** At the ventral position the density of CR^+^ neurons is reduced in some, but not all mice. **G)** In the dorsal contralateral subiculum, the density of CR-expressing cells is comparable to control. **H)** At the level of the injection site, CR^+^ cells are reduced also on the contralateral side. **I)** Further ventral CR immunolabeling is comparable to the ipsilateral side with reduced density in some mice. **J-L)** Quantification of CR-expressing cells in the subiculum of NaCl-injected mice, ipsi- and contralateral subiculum of KA-injected mice 21 days after injection. Individual animals are color-coded, bars show means (cells/mm^2^) with standard error of the mean (SEM). **J)** At the dorsal position the means of the three tested groups do not differ (one-way ANOVA: p = 0.20). **K)** At the level of the injection site, the density of CR-expressing cells is reduced at the ipsilateral and contralateral side compared to control (one-way ANOVA: p = 0.007; Tukey’s multiple comparisons test: NaCl-KAi p = 0.02, NaCl-KAc p = 0.009). **L)** At the ventral position the quantification does not reveal any significant differences, but large variability across mice (one-way ANOVA: p = 0.17). Scale bars: 250 μm. *p<0.05, **p<0.01. CA1 *cornu ammonis* 1, d deep pyramidal cell layer, m middle pyramidal cell layer, s superficial pyramidal cell layer.

At 21 days after ihKA we did not observe any major changes in the dorsal subiculum of the ipsi- or contralateral hemisphere, respectively (Fig. 5D,G). In contrast, at the level of the injection site CR-expressing cells were more sparse and their processes appeared reduced in staining intensity and density in the ipsilateral (Fig. 5E) and contralateral subiculum (Fig. 5H). In ventral parts, CR^+^ cells were reduced in some mice on both sides (Fig. 5F, I) whereas others showed normal distribution of CR^+^ INs. Quantification confirmed a significant reduction of CR-expressing INs at the level of the injection site for the ipsi- and contralateral subiculum (Fig. 5K), but not at the other positions (Fig. 5J,L).

Cells expressing CB(n=5) were primarily located in the superficial and middle layers of the proximal subiculum at the dorsal and the injection site level in controls (Fig. 6A, B). These densely positioned cells of low staining intensity likely reflect CB^+^ principal cells, as previously shown^32^, whereas some cells with higher staining intensity, scattered across all layers, represent putative INs. In ventral sections, the scattered cells were also visible in the distal subiculum, but the densely positioned cells were confined to proximal parts (Fig. 6C).

**Figure 6:**
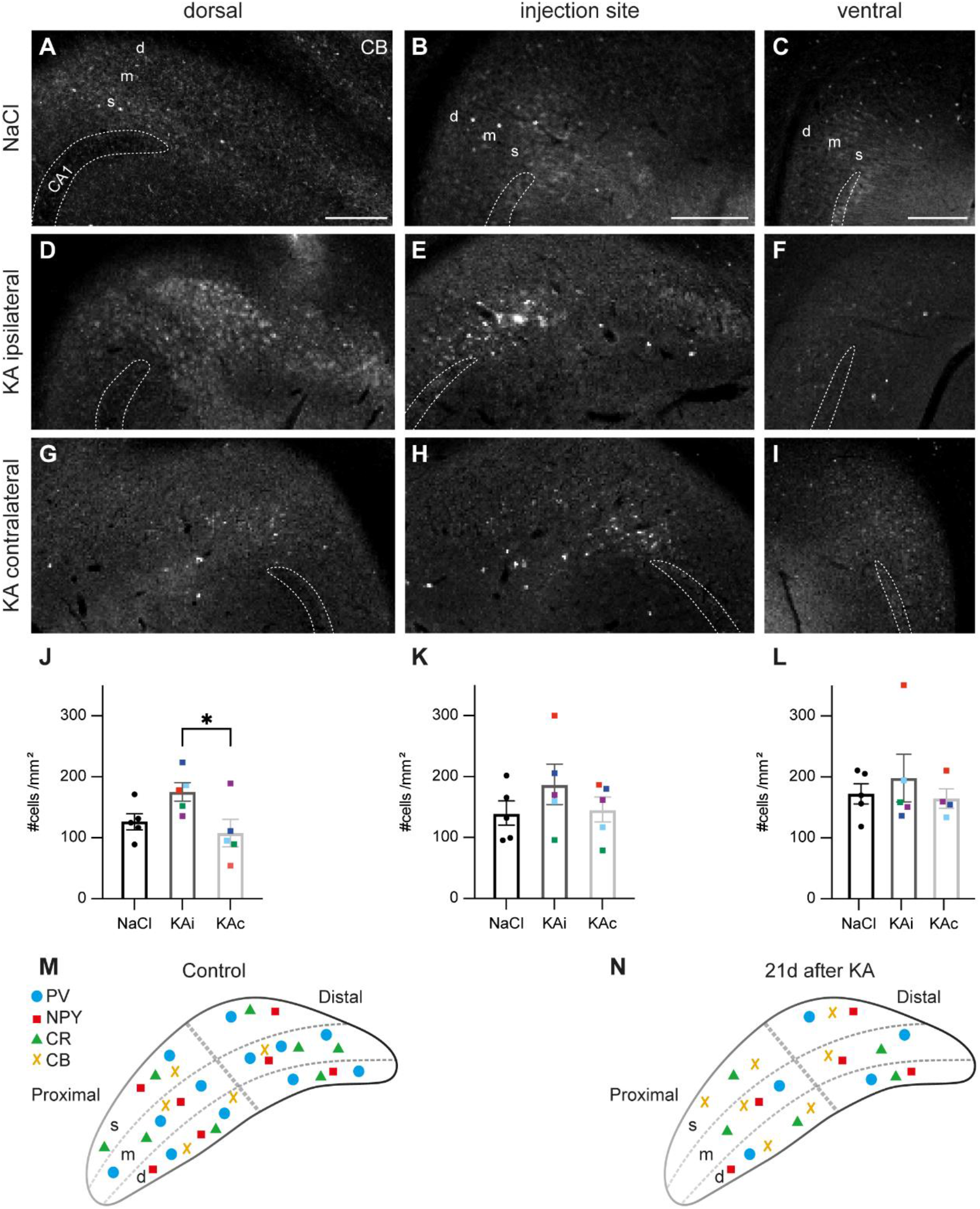
Cell density of calbindin-labeled cells is mildly increased under epileptic conditions. Representative images immunolabeled for calbindin (CB) from the subiculum of control (**A-C**) and epileptic mice (**D-I**). Borders of the CA1 pyramidal cell layer are indicated in all panels. **A-B)** In the dorsal subiculum and at the level of the injection site of NaCl-injected mice CB-expressing cells are mainly located in the superficial (s) and middle (m) pyramidal cell layer of the proximal subiculum. **C)** Ventrally, CB^+^ cells extend into the distal part of the subiculum. **D)** At 21 days after KA, staining intensity in the dorsal ipsilateral subiculum is increased compared to control. Cells expressing CB appear more clustered and malformed. **E)** At the level of the injection site and further ventral (**F**) the immunolabeling is slightly increased compared to control. **G)** In the contralateral dorsal subiculum CB staining appears weaker than in controls. **H)** On the level of the injection site and further ventral (**I**) distribution and intensity of labeled cells is comparable to controls. **J-L)** Quantification of CB-expressing cells in the subiculum of NaCl-injected mice, ipsi- and contralateral subiculum of KA-injected mice 21 days after injection. Individual animals are color-coded, bars show means (cells/mm^2^) with standard error of the mean (SEM). **J)** At the dorsal position an increase in mean density of CB-expressing cells was measured for the ipsilateral compared to the contralateral subiculum (one-way ANOVA p=0.05, Tukey’s multiple comparisons test KAi-KAc p = 0.04). **K)** At the injection site density was comparable for control and KA (one-way ANOVA: p = 0.39). **L)** Same for the ventral subiculum (one-way ANOVA: p = 0.68). Scale bars: 250 μm. *p<0.05. CA1 *cornu ammonis* 1, d deep pyramidal cell layer, m middle pyramidal cell layer, s superficial pyramidal cell layer. **M)** Schematic illustration of placement of subicular interneurons under control (left) versus epileptic conditions 21 days after KA (right) in all layers of the ipsilateral subiculum. Individual interneuron types are given in the legend.

In epileptic mice (n=5), a strong upregulation of CB staining was visible mainly in superficial and middle layers of the proximal but also in the distal ipsilateral subiculum at all DV levels (Fig. 6D-F). Labeling was often diffuse and cells appeared clustered and malformed, in particular close to CA1 (Fig. 6D,E). In contrast, CB staining in the contralateral subiculum was comparable to controls (Fig. 6G-I). The diffuse labeling impeded the quantification of CB^+^ cells which is reflected in high variability, already in controls, but even more after ihKA injection in both hemispheres (Fig. 6J-L). Accordingly, no differences in cell numbers could be detected except between the dorsal ipsi- and contralateral subiculum (Fig. 6J).

## Discussion

Our study provides a comprehensive insight into the pathological changes of the subiculum in a focal MTLE model with hippocampal sclerosis. Degenerating neurons in the ipsilateral subiculum were detected already 2 days after ihKA and neuronal density was persistently lower in chronic epilepsy. Specifically, we found the loss of GABAergic cells, mainly of PV^+^ INs and to a lesser extent of NPY^+^ and CR^+^ INs. In contrast, the upregulation of NPY and CB, in particular in non-GABAergic cells, suggests compensatory processes. Together, our data show that in the ihKA mouse model, the subiculum is involved in epileptic activity and undergoes epilepsy-associated histological changes of similar nature but less pronounced than in the hippocampus.

The degeneration of neurons early after ihKA injection is in agreement with the hypothesis that cell death in the subiculum contributes to epileptogenesis in the network beyond the hippocampus. Remarkably, neuronal degeneration was more pronounced at the dorsal position than at the level of the injection site, which might be due to the horizontal slicing direction which at the most dorsal levels does not cut exactly perpendicular to the longitudinal axis of the hippocampal formation. In addition, the axonal projections from the hippocampus to the subiculum span a considerable septo-temporal extent^33^, which might result in larger areas affected. With increasing distance from the injection site, the subiculum was better preserved in agreement with our previous studies investigating the hippocampus^23,25,26^. Interestingly, neuronal loss was more pronounced in the proximal than in the distal subiculum at the injection site and further ventral. Although this might be due to the mere proximity to the KA injection site, similar results have previously been found in animal models with (positionindependent) systemic chemoconvulsant injections in rats and mice^12,34^. In addition, the superficial and middle layers were more affected than the deep layer, suggesting a positiondependent differential vulnerability. Indeed, expression of molecular markers, intrinsic firing patterns and in- and outward connectivity differ between pyramidal cells in the proximo-distal as well as the transverse axis of the subiculum^35,36^, which might render them differentially vulnerable. A similar position-dependent diversity of neuronal properties is also conceivable among IN populations.

There is less mapping data available concerning the distribution of IN populations in the subiculum than for the hippocampus. We therefore chose *Gad67* as a general IN marker and four markers of specific populations to map their distribution in controls under our experimental conditions (Fig. 6M). Cells expressing PV were the largest fraction with somata in all layers and processes extending into the molecular layer. We found PV^+^ INs distributed nearly equally in the proximal and distal subiculum, which is in accordance with previous work^32^. Cells expressing NPY, of which most were GABAergic in controls, showed a gradient from dorsal to ventral and were more numerous in the proximal than in the distal subiculum. CR^+^ cells were evenly distributed at low density across the whole pyramidal layer of the proximal and distal subiculum. CB is not only a marker for INs, but also for subicular principal cell populations^31,32^. From their shape and location, more faintly stained cells represent principal cells, whereas strongly stained cells represent INs. In dorsal sections both CB^+^ cell types were mostly confined to the proximal subiculum but at further ventral levels the faintly stained cells were restricted to the proximal, whereas strongly stained cells were scattered throughout the subiculum. In the present study, we did not map INs expressing somatostatin (SOM), cholecystokinin (CCK) or vasoactive intestinal peptide (VIP), but this will also be of relevance in the future.

In epileptic mice, PV^+^ IN were lost by >40% in the dorsal subiculum and reduced at al DV positions which is in agreement with data from systemic rodent MTLE models^34,37,38^ and patients^9^. In accordance with the unilateral injection, there was no major loss of PV-expressing INs in the contralateral subiculum; occasional stronger contralateral PV staining might result from stronger somatic PV expression, but also from compensatory sprouting of PV^+^ terminals similar to the contralateral hippocampus^39^.

Double-labeling for NPY and *Gad67* mRNA revealed that these INs were also vulnerable to the ihKA injection, albeit less than PV^+^ INs in agreement with observations in the hippocampus^23^. For CR^+^ INs we found a reduction at the injection site whereas at the dorsal site an upregulation of CR in different cell types might have masked the reduction of CR^+^ INs. Generally, controversial data exist for the human subiculum in MTLE ranging from a reduction^40^ to normal density^41^, and high variability was also evident in systemic rat models^37,38^. The heterogeneous nature of CR-expressing neurons might play a role and require more elaborate investigation of subpopulation vulnerability. As mentioned earlier, we did not analyze populations expressing SOM, VIP or CCK, but the strong loss of *Gad67* mRNA-expressing cells in the dorsal subiculum suggests also reductions for these populations.

How might the reduced IN density affect population activity in the subiculum? It has been shown that during SWRs in the LFP subicular pyramidal cells either increase or decrease their firing and that these divergent cell types correspond to bursting and regular firing neuron types, respectively^42^. Interestingly, regular firing cells receive stronger synaptic inhibition during SWRs, in particular from PV^+^ INs which themselves are strongly activated during SWRs. Given the reduction in PV^+^ (and other) INs after ihKA injection, a shift towards more cells in bursting mode and hence from physiological SWRs to pathologic epileptic bursts is conceivable. In addition, despite the loss of CA1 input to the subiculum, projections from CA2 persist (own unpublished observations) and inputs from the entorhinal cortex, which strongly target the subiculum^43,44^ are also very likely to persist as we have shown it for the DG^13^. In coincidence with reduced feed-forward inhibition these excitatory projections might strongly activate the subiculum.

Such high excitatory pressure might also cause compensatory mechanisms to restore network balance. Indeed, we observed the upregulation of NPY after ihKA, comparable to epileptic rats^45^. This did not only affect the ipsilateral, but also the contralateral subiculum, which presented only little cell loss. Various *in vitro* and *in vivo* studies in animal models^46,47^, as well as in human tissue^48,49^ have shown NPY acting as endogenous anti-convulsant in the hippocampal network. Determining whether NPY plays a role in seizure suppression in the subiculum would require experiments with knock-out mice for NPY or its receptors.

The role of CB in epilepsy is heavily discussed: some studies claim the neuroprotective role of intracellular CB via its Ca^2+^-buffering potential, whereas others indicate that a reduction of CB, as observed in human and rodent granule cells in MTLE, might be beneficial via increasing Ca^2+^-dependent inactivation of Ca^2+^-channels thereby reducing Ca^2+^-influx^50^. Based on the current data, we do not dare to speculate about whether the CB upregulation in the subiculum in our model plays a neuroprotective role or leads to a greater disposition of the cells to hyperactivate.

## Conclusion

In summary, our data point towards an important role of the subiculum in the epileptic network by undergoing a 50% loss of INs and altered expression of NPY and Ca^2+^-binding proteins. This might render the subiculum into a hub which is disinhibited enough, and on the other hand presents sufficient neuronal preservation to contribute to the generation and/or the propagation of epileptic activity from the hippocampus to other parts of the brain.

## Acknowledgements

This work was funded by the Research Commission of the Medical Faculty, University of Freiburg (HAE2149/20 to UH), the German Research Foundation (HA 1443/11-1 to CAH), the BrainLinks-BrainTools Center, which is funded by the Federal Ministry of Economics, Science and Arts of Baden-Württemberg within the sustainability program for projects of the ExcellenceInitiative II. NB received a fellowship from the Center for Basics in Neuromodulation, University of Freiburg.

## Author contributions

Conceptualization: UH, ST; Data Curation: UH; Formal Analysis: JF, NB, UH; Funding Acquisition: UH, CAH; Investigation: JF, NB, ST, UH; Supervision: UH; Visualization: NB, UH; Writing -Original Draft Preparation: JF, NB, UH; Writing, Review & Editing: CAH, ST.

## Conflict of interest / Ethical publication statement

None of the authors has any conflict of interest to disclose.

We confirm that we have read the Journal’s position on issues involved in ethical publication and affirm that this report is consistent with those guidelines.

## Notes

### Competing Interest Statement

The authors have declared no competing interest.

